# ProtAnnot: an App for Integrated Genome Browser to display how alternative splicing and transcription affect proteins

**DOI:** 10.1101/025924

**Authors:** Tarun Mall, John Eckstein, David Norris, Hiral Vora, Nowlan Freese, Ann E. Loraine

## Abstract

**Summary:** One gene can produce multiple transcript variants encoding proteins with different functions. To facilitate visual analysis of transcript variants, we developed ProtAnnot, which shows protein annotations in the context of genomic sequence. ProtAnnot searches InterPro and displays profile matches (protein annotations) alongside gene models, exposing how alternative promoters, splicing, and 3’ end processing add, remove, or remodel functional motifs. To draw attention to these effects, ProtAnnot color-codes exons by frame and displays a cityscape graphic summarizing exonic sequence at each position. These techniques make visual analysis of alternative transcripts faster and more convenient for biologists.

**Availability and Implementation:** ProtAnnot is a plug-in App for Integrated Genome Browser, an open source desktop genome browser available from http://www.bioviz.org.

## 1 INTRODUCTION

Many genes produce multiple transcript variants due to alternative splicing, alternative promoters, and alternative 3’ end processing. Often these transcript variants encode proteins with different amino acid sequences and thus different functions. We and other groups have often used protein annotation methods to detect when this occurs. For example, we used BLOCKS, InterPro, and TMHMM to show that alternative transcription frequently remodels or deletes conserved regions and trans-membrane spans in human and mouse proteins (Cline, et al., 2004; Loraine, et al., 2002).

However, even now it is difficult for biologists to perform similar analysis. Using Web tools (Rodriguez, et al., 2015), biologists can identify conserved regions in proteins encoded by different splice variants, but mapping those regions back onto gene structures is time-consuming and error-prone.

To address this, we developed ProtAnnot as a new plug-in extension for the Integrated Genome Browser (IGB). IGB is a highly interactive, desktop genome browser that helps biologists explore and analyze experimental data from genomics, especially RNA-Seq data (Nicol, et al., 2009). Using ProtAnnot together with IGB, users can achieve deeper insight into how alternative transcription affects protein sequence and function.

### 2 RESULTS

ProtAnnot enables fast, efficient visual analysis of the impact of alternative transcription on proteins by extending standard genome browser iconography, in which linked blocks represent transcript structures and block thickness indicate translated regions.

ProtAnnot improves on this in three ways. First, it uses exon fill colors to show the frame of translation, revealing frame shifts across transcript variants. By comparing exon colors between transcripts, a user can quickly determine if they encode the same protein without having to zoom in to see the amino acid sequence.

Second, ProtAnnot introduces an exon summary graphic, a series of blocks at the bottom of the display whose heights indicate the number of exons overlapping each position. Height differences between blocks signal where models differ. By scanning the exon summary, users can easily identify difference regions (English, et al., 2010), sequences that are differentially included in transcripts due to alternative splicing, promoters, or 3'-end processing. The exon summary graphic draws attention to these regions by exploiting our native ability to notice discontinuities in a horizon (Fig. 1).

**Fig. 1.**
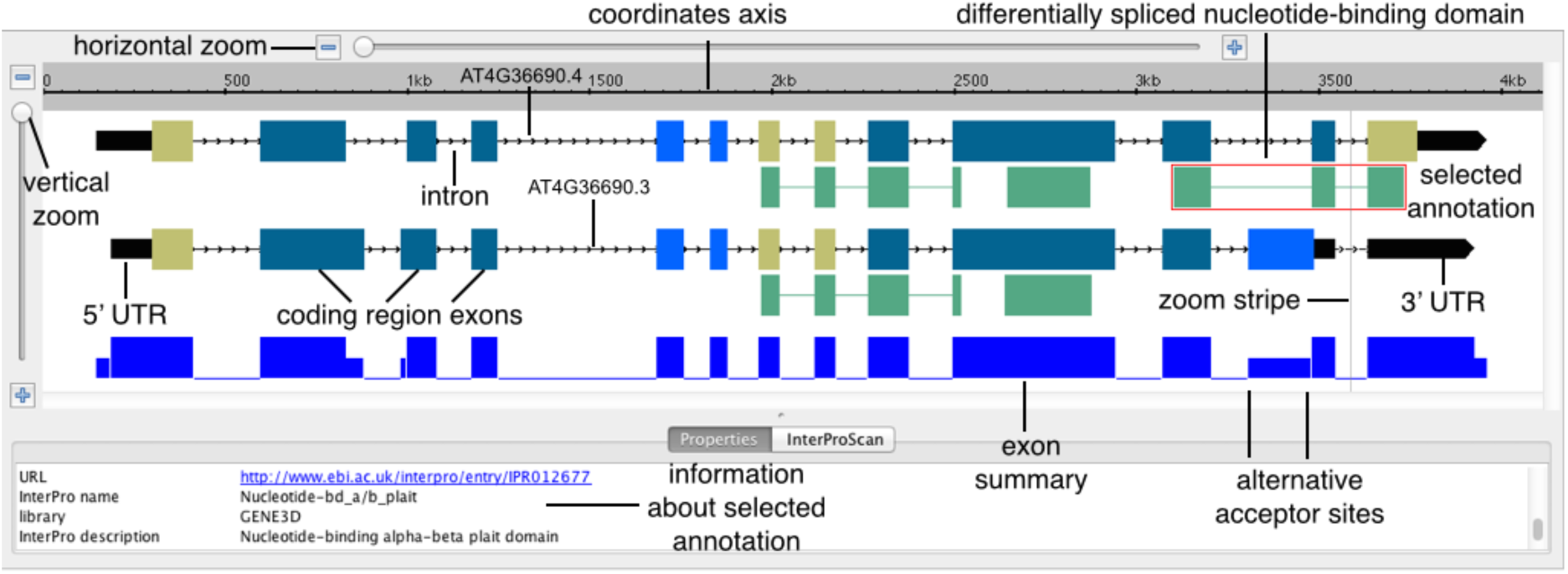
ProtAnnot visualization of *Arabidopsis thaliana* gene AT4G36690 encoding splicing regulator U2AF65 shows how alternative splicing deletes a nucleotide-binding domain. Different colors for coding region exons indicate different frames of translation.

Third, ProtAnnot exposes how different regions of a gene may encode different functions by displaying protein annotations next to their respective transcripts. In ProtAnnot, these protein annotations appear as single-or multi-span linked blocks beneath the transcripts that encode them. A thin line links spans from the same motif; note that these matches often span introns.

To use ProtAnnot, users select ProtAnnot in the IGB App Manager (available from the Plug-ins tab in IGB 8.5); this downloads ProtAnnot to a local cache. A new menu item labeled “Start ProtAnnot” then appears in the IGB Tools menu. Next, the user selects one or more gene models within the IGB main display window and selects “Tools > Start ProtAnnot”. This opens ProtAnnot in a new window, which shows the selected gene models with color-coded exons and exon summary graphic.

To search InterPro using ProtAnnot, users select the InterProScan tabbed panel and click a button labeled “Run InterProScan,” which opens a new window search options available from the InterProScan Web service hosted at the European Bioinformatics Institute. Users then select one or many databases to search, enter an email address, and run the search. Note that the InterProScan Web service maintainers require the user’s contact information to run the search service.

When the search finishes, ProtAnnot adds newly found protein annotations to the display, below their respective transcripts. ProtAnnot also updates the status message with a link to an XML file hosted on the EBI Web site containing the “raw” results; this is mainly a convenience for developers.

Clicking a protein annotation opens the Properties tab, which lists information about that particular profile or motif. Depending on the selected item, this can include its InterPro identifier, name, description, and a link to the InterPro Web site. Users can also shift-click to select multiple annotations, putting all of the available information side-by-side, allowing for direct comparison.

Figure 1 shows an example visualization that highlights how ProtAnnot can lead to new discoveries. This example shows a gene from the model plant *Arabidopsis thaliana* encoding a homolog of U2AF65, part of the U2AF dimer that recruits the U2 snRNP complex to the branchpoint adenosine residue during the early steps of splicing. Coding region exons are color-coded to indicate frame of translation, making it easy to notice when overlapping exons encode the same or different peptides. Likewise, the exon summary graphic highlights differences in splicing between gene models.

Previously, we analyzed RNA-Seq data from pollen and leaves and found that in pollen, AT4G36690.3 is the dominant isoform, but in leaves, AT4G36690.4 predominates (Loraine, et al., 2013). As shown in Figure 1, visualizing the gene models in ProtAnnot exposes potential functional consequences of pollen-specific splicing of U2AF65A. The pollen-specific isoform (lower) lacks a region encoding a nucleotide-binding alpha-beta domain, which is involved in RNA-binding. Thus, alternative splicing of U2AF65A is likely to have important functional consequences.

As with IGB, we developed ProtAnnot using the GenoViz SDK, a Java toolkit for building genome browsers (Helt, et al., 2009). By using GenoViz, we were able to implement advanced visualization techniques familiar to IGB users with minimal effort. These include: user-settable zoom focus indicated by a zoom stripe graphic, fast animated zooming, edge matching of selected items, and selectable Glyphs.

Search results can be saved and reopened later, which saves time for the user and also reduces load on the InterProScan Web service. ProtAnnot saves results to an XML format file (extension “.paxml”) that contains a slice of genomic sequence surrounding the gene models, an offset indicating the relationship between the slice and the larger reference, transcript structures using coordinates relative to the slice, and protein annotations in protein sequence coordinates.

## 3 CONCLUSION

ProtAnnot benefits users by exposing how gene structures affect protein sequence and function. As such, ProtAnnot complements the MI Bundle, another IGB extension that links genomic features to protein interaction and structure viewers (Céol and Müller, 2015). Like MI Bundle, ProtAnnot highlights relationships between the language of DNA (exons, introns, codons) and the more structure-oriented language of protein sequence, thus helping biologists achieve deeper understanding of gene function.

## ACKNOWLEDGEMENTS

John Nicol and Gregg Helt wrote early versions of ProtAnnot. This work was supported by National Institutes of Health [grant number R01GM103463].

